# Altered retinal structure and function in Spinocerebellar ataxia type 3

**DOI:** 10.1101/2022.01.10.475670

**Authors:** Vasileios Toulis, Ricardo Casaroli-Marano, Anna Camós-Carreras, Marc Figueras-Roca, Bernardo Sánchez-Dalmau, Esteban Muñoz, Naila S. Ashraf, Ana F. Ferreira, Naheed Khan, Gemma Marfany, Maria do Carmo Costa

## Abstract

Spinocerebellar ataxia type 3 is an autosomal dominant neurodegenerative disorder caused by expansion of a polyglutamine (polyQ)-encoding CAG repeat in the *ATXN3* gene. Because the ATXN3 protein regulates photoreceptor ciliogenesis and phagocytosis, we aimed to explore whether expanded polyQ ATXN3 impacts retinal function and integrity in SCA3 patients and transgenic mice.

We evaluated the retinal structure and function in five patients with Spinocerebellar ataxia type 3 and in a transgenic mouse model of this disease (YACMJD84.2, Q84) using, respectively, optical coherence tomography (OCT) and electroretinogram (ERG). We further determined in the transgenic mice: a) the retinal expression pattern of ATXN3 and assessed the distribution of cones and rods by immunofluorescence (IF); and b) the retinal ultrastructure by transmission electron microscopy (TEM).

Some patients with Spinocerebellar ataxia type 3 in our cohort revealed: i) reduced central macular thickness indirectly correlated with disease duration; ii) decreased thickness of the macula and the ganglion cell layer, and reduced macula volume inversely correlated with disease severity (SARA score); and iii) electrophysiological dysfunction of cones, rods, and inner retinal cells. Transgenic mice replicated the human OCT and ERG findings with aged homozygous Q84/Q84 mice showing a stronger phenotype accompanied by further thinning of the outer nuclear layer and photoreceptor layer and highly reduced cone and rod activities, thus supporting severe retinal dysfunction in these mice. In addition, Q84 mice showed progressive accumulation of ATXN3-positive aggregates throughout several retinal layers and depletion of cones alongside the disease course. TEM analysis of aged Q84/Q84 mouse retinas supported the IF ATXN3 aggregation findings by revealing the presence of high number of negative electron dense puncta in ganglion cells, inner plexiform and inner nuclear layers, and further showed thinning of the outer plexiform layer, thickening of the retinal pigment epithelium and elongation of apical microvilli.

Our results indicate that retinal alterations detected by non-invasive eye examination using OCT and ERG could represent a biological marker of disease progression and severity in patients with Spinocerebellar ataxia type 3.

## INTRODUCTION

Spinocerebellar ataxia type 3 (SCA3), also known as Machado-Joseph disease (MJD) (OMIM#109150), is a late onset autosomal dominant neurodegenerative disorder and the most common inherited ataxia in the world^1^. Spinocerebellar ataxia type 3 is caused by an expanded CAG triplet repeat size in the *ATXN3* gene^2^, encoding a polyglutamine (polyQ) tract in the ATXN3 deubiquitinating enzyme. In healthy individuals, *ATXN3* harbors 12 to about 51 CAGs, whereas in patients with Spinocerebellar ataxia type 3 the gene contains an expanded (CAG)_n_ tract that ranges from ~60 to 87 triplets^3,4^. Intermediate length alleles (45-60 CAG repeats) show incomplete penetrance or differential clinical presentation^5^. The length of the *ATXN3* expanded CAG repeat correlates directly with phenotypic severity and disease progression rate, and indirectly with the age at onset of the first clinical manifestations^6–11^.

As a deubiquitinating enzyme, wild type ATXN3 modulates proteotoxic stress, aging, and cell differentiation^12^. Perturbation of ATXN3 native conformation by a pathogenic expanded polyQ tract alters its interaction with other protein partners and favors the formation of insoluble aggregates inducing cell stress and leading to eventual cell death^12–15^. Patients with Spinocerebellar ataxia type 3 show ATXN3 aggregates and selective cell loss in the cerebellum, brain stem, mid brain, spinal cord, and peripheral nerves^11,15–17^. These central nervous system (CNS) damages are reflected by progressive ataxia, ophthalmoplegia, dysarthria, dysphagia, pyramidal signs (rigidity and spasticity), dystonia, amyotrophy, and visual symptoms^13,15,18,19^.

Although neurodegeneration in dominantly inherited ataxias is usually cell-specific and mostly restricted to the CNS, patients with Spinocerebellar ataxia type 7 also show progressive retinal neurodegeneration with photoreceptor loss and retinal pigment epithelium (RPE) lesions^20–22^. Because the clinical manifestations reflecting brain degeneration in Spinocerebellar ataxia type 7 are very similar to those in Spinocerebellar ataxias type 1, 2 and 3^15^, a certain degree of retinal degeneration/dysfunction might also be associated to these polyQ spinocerebellar ataxias^23^. While the retina is part of the CNS, it is more amenable than the brain to morphological and physiological observation. In fact, the eye may be considered a window to the brain, since the retina may display very early symptoms of neurological and psychiatric disorders^24^.

Identification of initial symptoms of spinocerebellar ataxias is extremely important as these signs can be tracked over the course of clinical trials enrolling pre-symptomatic or early-stage disease patients. Visual disturbance, such as double vision, has been described as the initial symptom, preceding gait ataxia in many years, in about 10% of patients with Spinocerebellar ataxia type 3^25,26^ and, more recently, abnormal eye movements have been confirmed in pre-ataxic carriers^27^ correlating with disease severity in patients^27, 28^. Whereas patients with Spinocerebellar ataxia type 3 show retinal nerve fiber layer (NFL), ganglion cell layer (GCL) and macular thinning^29–31^, there is no reported evidence of decreased activity of retinal cells in these patients. Because ATXN3 is highly expressed in the retina^32^ and plays a role in retinal ciliogenesis and RPE phagocytosis^33^, here we aimed to explore whether expanded polyQ ATXN3 impacts retinal function and integrity in Spinocerebellar ataxia type 3 patients and transgenic mice.

## MATERIALS AND METHODS

### Ethics statement

All procedures were performed according to the ARVO statement for the Use of Animals in Ophthalmic and Vision Research, as well as the regulations of the Animal Care facilities at the University of Michigan. All animal procedures were evaluated and approved by the Animal Research Ethics Committee of the University of Michigan Committee on the Use and Care of Animals (animal protocols PRO00008371, and PRO00008409).

Regarding the human patients, this study followed the ethical precepts of the Declaration of Helsinki (Fortaleza, Brazil, Oct 2013) and was approved by the hospital ethics committee (CEIC, Hospital Clinic de Barcelona; Ref HCB/2019/0794).

### Spinocerebellar ataxia type 3 patients and clinical assessment

All clinical assays were carried out under patient informed consent. The use of personal data complied with local regulations (Spanish Law 05/2018). In this study, patients were evaluated by specialists in neurology, neuro-ophthalmology and ocular electrophysiology. The neurological examination obeyed the standard criteria for pathology and the cases were duly classified according to SARA (Scale for the Assessment and Rating of Ataxia) - ranges from 0 (no ataxia) to 40 (most severe ataxia) - which is currently one of the instruments of choice to evaluate the grade of disease severity^34,35^. SARA scale was used as primary outcome to provide preliminary clinical evidence – early and late clinical disease - for further research. Patients were also classified according to disease stage as follows^36^: stage 0, no gait difficulties; stage 1, disease onset, as defined by onset of gait difficulties; stage 2, loss of independent gait, as defined by permanent use of a walking aid or reliance on a supporting arm; stage 3, confinement to wheelchair, as defined by permanent use of a wheelchair. Genetic diagnosis had previously been carried out in the setting of their clinical care (Genetic Service of Hospital Clinic de Barcelona) following standard procedures. Spinocerebellar ataxia type 3 diagnosis was definitively established when a genetic study confirmed the presence of a CAG trinucleotide repeat expansion with more than 60 repeats in the *ATXN3* gene on chromosome 14q24.3-q31^37^.

### Optical coherence tomography (OCT) and ocular electrophysiological test in Spinocerebellar ataxia type 3 patients

Complete standard ophthalmologic examination was carried out for each participant, including best-corrected visual acuity (BCVA) with Snellen chart, intraocular pressure, anterior and posterior segment biomicroscopy, and ocular motility. Spectral-domain optical coherence tomography (OCT, AngioPlexTM CIRRUSTM HD-OCT 5000, Carl Zeiss Meditec, Inc., Dublin, USA) was performed to measure retinal parameters: central retinal, GCL and NFL thicknesses, and macular volume. OCT acquisition protocols were Optic Disc Cube (6×6 mm; 200×200 scans) and Macular Cube (6×6 mm; 521×128 scans) according to the OSCAR-IB and APOSTEL^38,39^ criteria. Image analysis was carried out following image segmentation procedure with the built-in manufacturer software, based on algorithms that compare each measurement with the internal database of the device, obtained by thousands of results in normal population, and calculate the deviation of normality. Eelectrophysiological testing (RETI-scan science, Roland-Consult, Brandenburg, Germany) by full-field flash electroretinogram (ffERG, overall retinal function exploration) and multifocal electroretinogram (mfERG, central retinal function exploration) was conducted according to the International Society for Clinical Electrophysiology of Vision (ISCEV) standards^40^.

### Animal procedures

Spinocerebellar ataxia type 3 YACMJD84.2 transgenic mice (C57BL/6 background) (from now on, Q84 mice)^41^ were housed in cages with a maximum number of five animals and maintained at the University of Michigan in a standard 12-hour light/dark cycle with food and water *ad libitum*. Mouse genotyping was performed using genomic DNA isolated from tail biopsies at the time of weaning, and genotypes of all studied litter mate mice - hemizygous transgenic (Q84), homozygous transgenic (Q84/Q84), and non-transgenic (WT) - were reconfirmed using DNA extracted from tails collected post-mortem. Total DNA was isolated using the DNeasy Blood and Tissue kit (Qiagen, Hilden, Germany), following the manufacturer’s instructions with minor modifications. Primers and PCR conditions for genotyping have been previously reported ^41^. Animals were anaesthetized with isoflurane and, once pedal withdrawal reflex was lost, mice were euthanized by decapitation. Eyes were immediately dissected from [6-8]-month-old and 1.5 year-old hemizygous Q84, homozygous Q84/Q84, and wild type (WT) littermate mice.

### Protein lysates and immunoblotting

Murine retinas were dissected and homogenized in RIPA lysis buffer (50 mM Tris pH 7.5, 1 mM EDTA, 150 mM NaCl, 0.5% NP40) containing protease inhibitors (Complete, Roche Diagnostics, Indianapolis, IN), followed by sonication and centrifugation. Total protein concentration in lysates (supernatants) was determined using the BCA method (Pierce™ BSA Protein Assay Kit, Thermo Fisher Scientific, Waltham, MA). Proteins (20-30 μg) were resolved in 10% SDS-PAGE gels, transferred onto PVDF membranes and blocked with 5% non-fat milk in TBS-T for 1 h, followed by an overnight incubation at 4°C with the primary antibodies rabbit anti-ATXN3 (anti-MJD 1:20000)^42^ or mouse anti-GAPDH (1:1000, ab9484 Abcam, Cambridge, UK), and subsequent incubation with the corresponding peroxidase-conjugated anti-rabbit or anti-mouse secondary antibodies (1:2000) for 1 h. Bands were visualized by treatment with the ECL-plus reagent (Western Lightning®, PerkinElmer, Waltham, MA) and exposure to autoradiography films.

### Immunofluorescent histochemistry (IHC)

Mouse eye cups were dissected, fixed, cryoprotected and sectioned for immunofluorescence detection as previously described^43^. For immunofluorescent histochemistry, 12 μm retinal sections were recovered on Superfrost Plus glass slides (Electron Microscopy Sciences, Hatfield, PA), dried 30-45 min at room temperature (RT), washed with PBS for 10 min, permeabilized with 0.5% Triton X-100 in PBS (20 min at RT), washed three times with PBS-T (0.1% Tween-20 in PBS) for 10 min, and blocked in 5% goat or sheep serum in PBS for 1 h. Primary antibodies were incubated overnight at 4°C diluted in blocking solution. The primary antibodies and corresponding dilutions were as follow - rabbit anti-ATXN3 (anti-MJD, 1:5000), mouse anti-rhodopsin (clone 1D4, 1:250, ab5414, Abcam, Cambridge, UK), rabbit anti-red/green opsin (1:300, AB5405, Merck Millipore, Burlington, MA), mouse antiacetylated α-tubulin (1:2000, T6793, Sigma Aldrich, San Luis, MO). Secondary antibodies Alexa Fluor 488 anti-mouse (1:300), Alexa Fluor 568 anti-rabbit (1:300), Lectin PNA conjugated to Alexa Fluor 488 (1:50) (all from Life Technologies, Grand Island, NY) and DAPI fluorescent die (1:300, Roche Diagnostics, Indianapolis, IN) were incubated at RT for 1 h. Finally, the slides were mounted in ProLong^®^ Gold (Invitrogen, Carlsbad, CA) and analyzed blindly for the genotype by confocal microscopy (SP2 or SP5, Leica Microsystems, Wetzlar, Germany, or Nikon A1, Tokyo, Japan).

### Whole mount retinal staining

Eyes from Q84/Q84 and WT littermate mice were enucleated, fixed and dissected as previously described^43^. Whole neural retinas with the photoreceptor layer upside were flattened by performing four symmetrical cuts on Superfrost Plus glass slides (Electron Microscopy Sciences, Hatfield, PA), fixed with 4% PFA for 1 h, washed three times in PBS for 10 min, and blocked for 1 h in 5% goat serum in PBS. The whole retinas were incubated at RT for 1 h with Lectin PNA conjugated to Alexa Fluor 647 (1:50, L32460, Life Technologies, Carlsbad, CA) in blocking solution, mounted in ProLong® Gold (Invitrogen, Carlsbad, CA) and analyzed by confocal microscopy (Nikon A1, Tokyo, Japan).

### Transmission Electron Microscopy (TEM)

Eyes from Q84/Q84 (N=3, 2F/1M) and WT (N=3, 1F/2M) mice were enucleated, the cornea was perforated using a needle to create a small hole, and eyes were immersed in fixative solution (2.5% glutaraldehyde, 2% PFA in 0.1 M cacodylate buffer) for 1 hour at RT, followed by an overnight incubation at 4°C. After several rinses with 0.1 M cacodylate buffer, eyes were post-fixed in 1% osmium tetroxide, rinsed in double distilled water to remove phosphate salt and then stained *en bloc* with aqueous 3% uranyl acetate for 1 hour. Eyes were dehydrated in ascending concentrations of ethanol, rinsed two times in propylene oxide, and embedded in epoxy resin. Eye blocks were sectioned into 70 nm ultra-thin sections and stained with uranyl acetate and lead citrate. Negative-stained sections were examined using a JEOL 1400 electron microscope at 80 kV. Images were recorded digitally and blindly to the genotype using a Hamamatsu ORCA-HR digital camera system operated using AMT software (Advanced Microscopy Techniques Corp., Danvers, MA).

### Mouse spectral domain-optical coherence tomography

*In vivo* images of mouse retinal structure were obtained with spectral domain-optical coherence tomography (SD-OCT) (Leica Microsystems, Buffalo Grove, IL). The mice received 1% tropicamide drops to stimulate eye dilation and were anesthetized with ketamine and xylazine (50 and 5 mg/kg body weight, respectively). Cube volumes consisting of 1000 A-scans by 100 B-scans over a 1.4 × 1.4-mm area centered on the optic nerve head (ONH) were taken for visualization of retinal anatomy. Retinal thickness was measured at 350 μm from the ONH using the Bioptigen Diver software. Measurements of thickness for different retinal layers were obtained in nasal, temporal, superior, and inferior regions of the retina. The four measurements were averaged to generate an average retinal thickness.

### Mouse electroretinogram

A Diagnosys Celeris System (Diagnosys, Lowell, MA) was used to assess mouse retinal function. After dark adaptation overnight, the mice received a drop of 0.5% tropicamide to stimulate eye dilation, a drop of 0.5% proparacaine to numb the eyes, and were anesthetized using an intraperitoneal injection of ketamine (50 mg/kg bodyweight) and xylazine (5 mg/kg bodyweight). A drop of 0.3% hypromellose (Genteal) was used to hydrate the corneas and to cushion the contact with the electrode stimulators. Scotopic responses were recorded for 0.01, 0.1, 1, 10, and 32 cd*s/m^2^ intensity stimuli. Mice were then light adapted for 10 minutes to a 30 cd/m^2^ white background light prior to photopic testing. Photopic responses were recorded sequentially using 10, 32, and 100 cd*s/m^2^ stimuli. Photopic flicker responses were then recorded using 20 cd*s/m^2^ cycled at 9.9 Hz. During testing, body temperature was maintained to 37°C by the built-in heating element.

### Statistical analyses

For statistical significance of data (for all assays, two eyes of n≥ 5 animals), equivalent standard deviation (SD) and normal distribution were first assessed using Bartlett and Shapiro-Wilk tests. When data followed a normal distribution and showed homogeneity of variance, two-way ANOVA or T-test were used for statistical significance analysis. For nonnormal and/or non-homogeneous variance distributions, non-parametrical Kruskal-Wallis one-way analysis or Mann-Whitney U-test were applied. and Analysis was performed using GraphPad Prism 7.03 (San Diego, CA) or Statgraphics Centurion XVII Statistics software (The Plains, VA). Spearman correlations and partial correlations with co-variant adjustments were performed using IBM SPSS Statistics for Windows, version 25.0 (IBM Corp., Armonk, N.Y., USA).

## RESULTS

### Patients with Spinocerebellar ataxia type 3 show mild to prominent retinal dysfunction

Because the retina is a neurological organ very sensitive to cellular stress and we recently uncovered that native ATXN3 is involved in retinal ciliogenesis and RPE phagocytosis^33^, we sought to evaluate whether the retina of patients with Spinocerebellar ataxia type 3 shows signs of mutant ATXN3-mediated cytotoxicity using non-invasive optical coherence tomography (OCT) and electroretinograms.

For this exploratory study, we reviewed the clinical records of 25 patients with confirmed clinical and genetic diagnoses for Spinocerebellar ataxia type 3. Among these, nine patients on disease stages 1 to 3^36^ were pre-selected for our study protocol, of which five patients (three females and two males) with a mean age of 59.6 years (range 45-83), in disease stages 1 (n=2; P1 and P2) and 3 (n=3; P3, P4 and P5) met the conditions for an ophthalmological and ocular electrophysiology evaluation (Supplementary Table 1). SARA scale^34,35^ was on average 13.5 [6-21.5], the mean CAG repeat number of the cohort was 68 [61-74], with an average disease duration of 14.2 years [8-23 years] (stage 1: mean 9 years and stage 3: mean 17.6 years) (Supplementary Table 1). BCVA was considered normal (range 20/30 - 20/20 in the Snellen scale) for the 10 eyes of the evaluated patients with Spinocerebellar ataxia type 3 (Supplementary Table 1).

A careful segmentation analysis of the OCT measurements revealed some specific alterations, such as macular, GCL and NFL peripapillary thinning, from mild to more pronounced, with a high index of symmetry (mean: 87.6%; 77% - 92%) for the thickness of the NFL (Figure 1, Supplementary Table 1). While the number of evaluated eyes of patients with Spinocerebellar ataxia type 3 was limited (n=10), we determined whether the central and average macular, GCL, and NFL thicknesses and macular volume were individually associated with the disease progression and severity by conducting correlation analyses of these measurements with the disease duration and SARA score. We first assessed if the expanded CAG repeat size and the patient’s age at the evaluation time have an effect in each individual OCT measurement. The expanded CAG size did not correlate with any evaluated parameters in our patients, but the central macular thickness inversely correlated with the age of our patients. Next, and adjusting for the age, we observed that the central macular thickness inversely correlated with disease duration (Figure 1B) but not with the SARA score (Figure 1C) suggesting that Spinocerebellar ataxia type 3 progression, but not severity, impact the central macular thickness in patients affected with this disease. We detected, however, that the average macular and the GCL thicknesses, and the macular volume inversely correlated with the SARA score (Figure 1C) further indicating that disease severity affects the macula and the GCL.

**Figure 1.**
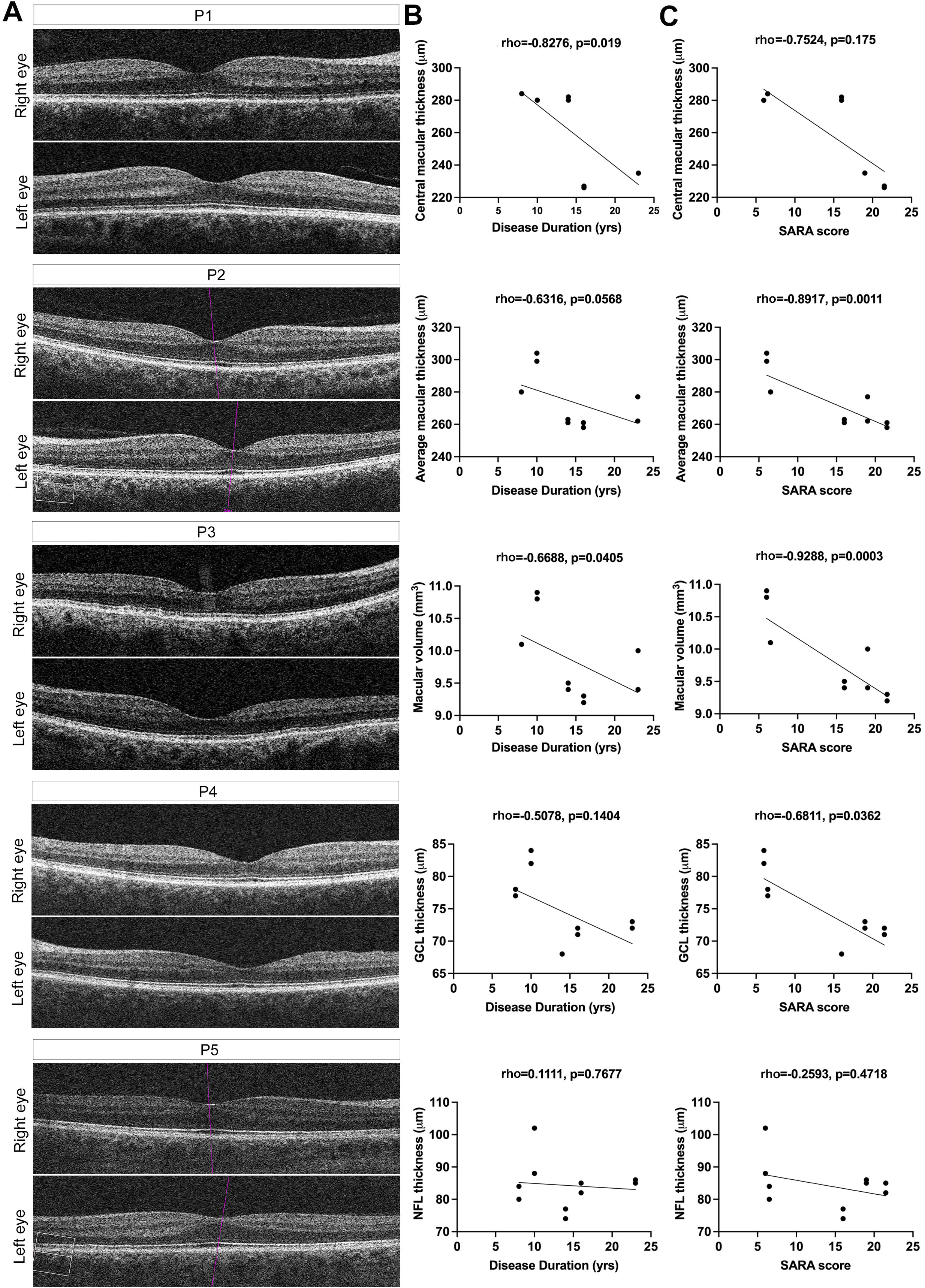
Patients with Spinocerebellar ataxia type 3 show progressive retinal thinning. **(A)** Optical coherence tomography (OCT) images of both eyes of patients with Spinocerebellar ataxia type 3 (P1 to P5). **(B)** Correlation of OCT measurements with disease duration revealing that central macular thickness and macular volume inversely associate with the disease duration. **(C)** Correlation of OCT assessments with the SARA score showing that average macular thickness, macular volume and ganglion cell layer (GCL) indirectly associate with the SARA score. Spearman correlation (rho) adjusted for age whenever needed with statistical significance set at p<0.05. NFL _ nerve fiber layer.

Multifocal electroretinogram (mfERG) assessment of patients with Spinocerebellar ataxia type 3 revealed impaired registers of the: a) macular peripheral rings in a stage 1 patient (P1) and in a stage 3 patient (P4) (Figure 2A, Supplementary Table 1); and b) both central and peripheral rings in another stage 3 patient (P3), thus indicating rod (peripheral) and cone (central) affectation in both stages 1 and 3 (Figure 2A, Supplementary Table 1). Patients P2 and P5 also showed mild alterations by mfERG, mostly in the cones of the fovea, and further evidence of peripheral alterations in patient P5. Concerning biological sex, interestingly the most affected patients were female (P1, P3 and P4). Importantly, correlation analyses of mfERG measurements with disease duration and SARA score (adjusted for age and the expanded CAG repeat size whenever needed) revealed that the response activity density (RAD) of the parafovea (ring 2) inversely correlated with the disease duration (Figure 2B), further indicating that the disease progression affects not only the central macula architecture (Figure 1B) but also its electrophysiological activity.

**Figure 2.**
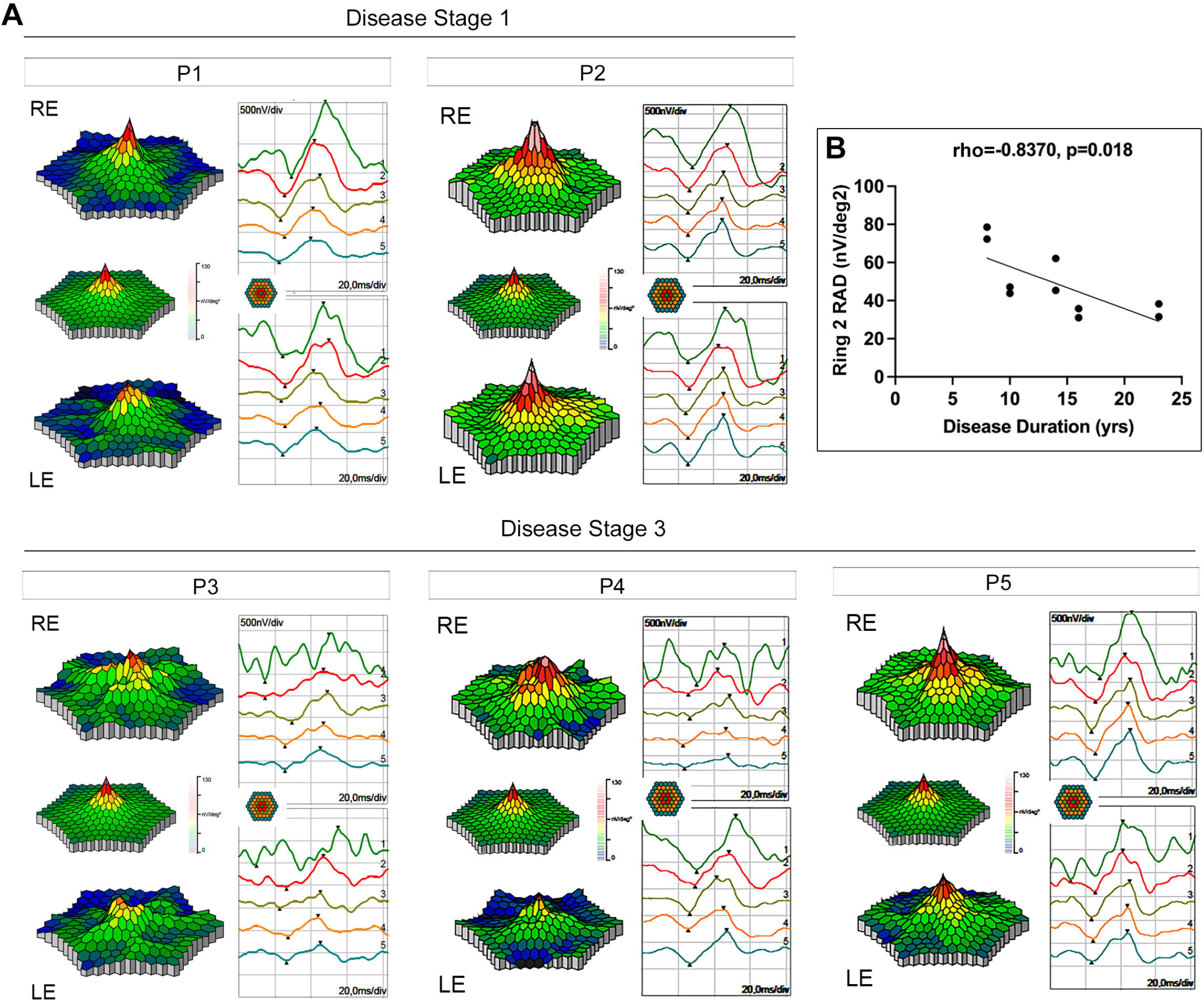
Multifocal ERG evaluation of patients with Spinocerebellar ataxia type 3 shows mild to severe retinal dysfunction and decreased activity of the parafovea with increased disease duration. (**A**) Precise topographical retinal function distribution and electrophysiological activity for rings 1 to 5 in patients with Spinocerebellar ataxia type 3. Peripheral activity reflecting rod activity shifts from green (functional) to blue (dysfunction), whereas cone-rich macula shifts from red to yellow/white. Normal mfERG is shown for comparison between the retinal geographic activity records of the right and left eye of each patient. Patients P1, P3 and P4 show a more severe dysfunction, in both peripheral and macular electrophysiological records, whereas P2 and P5 show a milder affectation. (**B**) The response amplitude density (RAD) of the parafovea inversely correlates with the disease duration. Statistical significance set by Spearman partial correlation (rho) adjusted for age at p<0.05. RE _ right eye; LE _ left eye; 1 _ R1, ring 1, fovea; 2 _ ring 2, parafovea; 3 _ ring 3, perifovea; 4 _ ring 4, near periphery; 5 _ ring 5, central part of the middle periphery; yrs, years.

Full-field flash electroretinogram (ffERG) evaluation of the same five above-described patients showed abnormal registers in one patient of stage 1 (P1) and in two patients of stage 3 (P3, P4), thereby revealing alterations in implicit times and amplitudes of a- and b-waves, in both photopic and scotopic conditions, respectively consistent with affectation of activity of cone and rod photoreceptors, and of ON-and OFF-bipolar and Müller cells (Supplementary Table 1). Moreover, correlation analyses of ffERG measurements with disease duration and SARA score revealed that: a) the implicit time of scotopic (dark-adapted) b-wave directly and inversely correlated with the disease duration, respectively, at low 0.01 cd.s/ms^2^ and medium 1.0 cd.s/ms^2^ flash strengths indicating that ON-bipolar cells respond slower to low stimulus and quicker to medium stimulus of light with the course of disease (Figure 3A); b) the implicit time of scotopic a-wave (10.0 cd.s/ms^2^) directly correlated with the disease duration and the SARA score revealing that rods respond slower with the disease progression and severity (Figure 3B); and c) the implicit time of oscillatory potentials (OPs) directly correlated with the disease duration implying that the interaction of inner retina cells (amacrine, bipolar and ganglion cells) slows down with the disease progression (Figure 3C).

**Figure 3.**
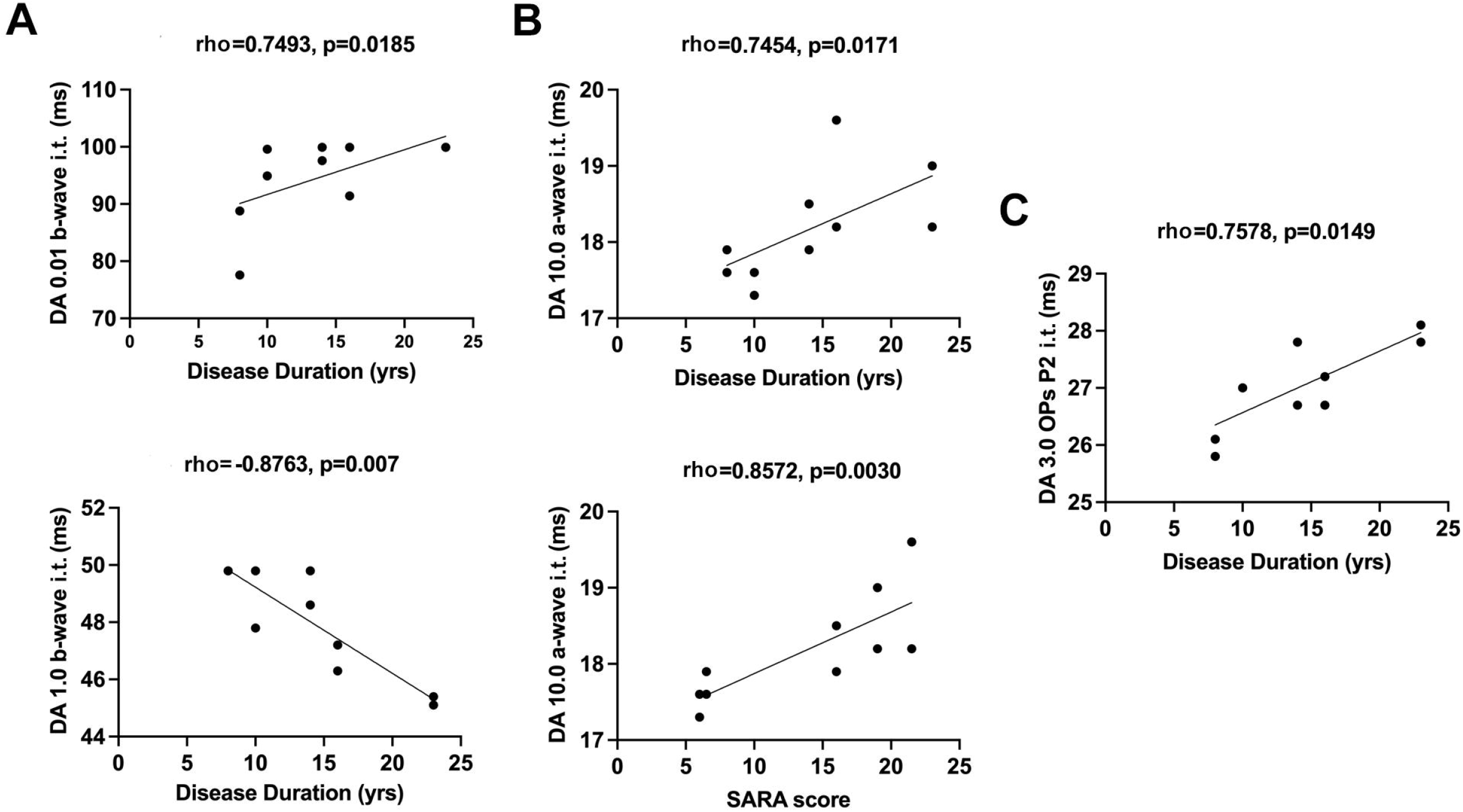
Response of retinal cells slows down with the duration and severity of Spinocerebellar ataxia type 3. (**A**) The implicit time (i.t.) of scotopic (dark-adapted, DA) b-wave directly and inversely correlates with the disease duration, respectively, at low 0.01 cd.s/ms^2^ (top panel) and medium 1.0 cd.s/ms^2^ (bottom panel) flash strengths. (**B**) The implicit time of scotopic a-wave (10.0 cd.s/ms^2^) directly correlates with the disease duration (top panel) and the SARA score (bottom panel). (**C**) the implicit time of oscillatory potentials (OPs) directly correlated with the disease duration. Statistical significance set by Spearman partial correlation (rho) (adjusted for age and the size of the expanded CAG repeat when needed) at p<0.05. yrs - years.

Overall, these *in vivo* complementary structural and functional tests in patients with Spinocerebellar ataxia type 3 show slight progressive retinal thinning and decreased or delayed ERG responses of photoreceptors and other retinal cells related with disease duration and severity. Given our small patients’ sample, further confirmation of these findings in a large cohort of patients is required for firmer conclusions.

### Spinocerebellar ataxia type 3 transgenic mice show thinning of several retinal layers

We next sought to evaluate whether Spinocerebellar ataxia type 3 YACMJD84.2 (Q84) transgenic mice replicate the visual deficits shown by patients with this disease. Q84 mice expressing the full-length human *ATXN3* disease gene harboring an expanded CAG repeat (CAG_84_, Q84) have been extensively used in several preclinical studies for Spinocerebellar ataxia type 3^41,44–50^. While hemizygous Q84 mice (Q84) show a mild motor phenotype of late onset, homozygous Q84 mice (Q84/Q84), with a higher overexpression of the pathogenic human allele, show motor deficits and low body weight beginning as early as 6 weeks of age^46^. Q84/Q84 male and female mice are equally affected with progressive phenotypes until early mortality in average around 50-60 weeks of age^46^. These deficits are accompanied by neuropathological features that recapitulate features of human Spinocerebellar ataxia type 3, including intranuclear accumulation of ATXN3 and ATXN3-positive aggregates in several brain regions known to be affected in patients^11,51,52^. As Q84 transgenic mice replicate many aspects of the human disease we investigated whether they also reproduce SCA3-like visual deficits. Because, to the best of our knowledge, there were no previous retinal studies in Q84 mice, we first confirmed by Western blot that the human *ATXN3* transgene is expressed in the retina. The human expanded-polyQ ATXN3 protein is clearly overexpressed in Q84/Q84 and Q84 mouse retinas compared to WT littermate controls (Supplementary Figure 1).

To evaluate potential effects on retinal anatomy, we used SD-OCT to measure the retinal layer thickness in both eyes of 10-12 month-old Q84 (N=5, 3F/32), Q84/Q84 (N=6, 2F/4M) and WT (N=7, 3F/4M) littermate mice (Figure 4A). Hemizygous Q84 retinas revealed a modest, but significant, reduction of 4% of total retinal layer thickness (Figure 4A,B) of controls due to thinning of the inner layer (Figure 4B), specifically of GCL and inner plexiform layer (IPL) (Figure 4C). On the other hand, total retinal layer thickness of homozygous Q84/Q84 mice was markedly decreased in 18% of that of WT mice (Figure 4A). More precisely, Q84/Q84 mice showed statistically significant thinning of both inner and outer retinal layers, with a respective reduction of 10% and 30% of controls (Figure 4B), which was consistent with decreased thickness in NFL, GCL, IPL, outer nuclear (ONL), and photoreceptor (PhR) layers (Figure 3C).

**Figure 4.**
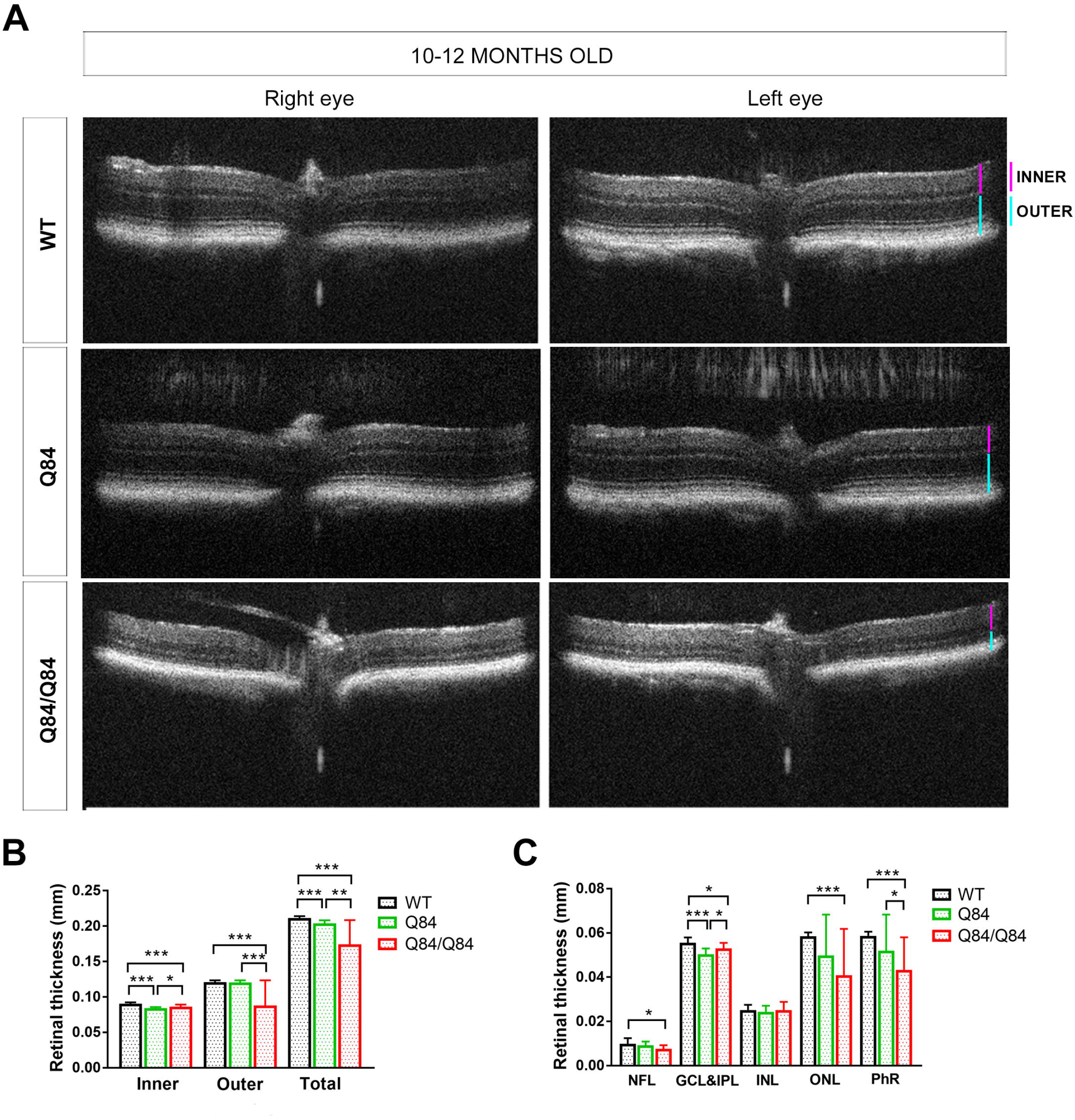
Optical coherence tomography reveals a general decrease in the thickness of the retinal layers in homozygous Q84/Q84 compared to wild-type mice. **(A)** Representative optical coherence tomography (OCT) images indicate thinner retinal layers (both in the inner and outer layers) of Q84/Q84 homozygous (n=6) compared to wildtype (WT) (n=7) mouse retinas. Hemizygous Q84 animals (n=5) show an intermediate phenotype. **(B, C)** Quantification of the retinal layer thickness observed in Q84 (hemizygous and homozygous) versus WT mice. Statistical significance, * p<0.05, ** p<0.01, *** p<0.001 (two eyes of 5-7 animals per group). PhR - photoreceptors, ONL - outer nuclear layer, INL - inner nuclear layer, IPL - inner plexiform layer, GCL - ganglion cell layer, NFL - nerve fiber layer.

Overall, both Q84 and Q84/Q84 retinas showed tightening of NFL and GCL reproducing the OCT findings in patients with Spinocerebellar ataxia type 3, in this and other studies^29–31^. Q84/Q84 retinas displayed further thinning of the outer layer (ONL and PhR layer) compared to controls, thus suggesting potential loss of photoreceptors in the transgenic retinas.

### Retinas of Spinocerebellar ataxia type 3 transgenic mice display electrophysiological deficits

To investigate whether retinal layer thinning in Q84 mice affected the electrophysiological response upon light-stimuli, we next conducted ERG testing in the same 10-12 month-old transgenic mice used for OCT assessment. We evaluated both cone (photopic) and rod (scotopic) responses in Q84 transgenic mice. The ERG response of hemizygous Q84 mice showed a normal cone response at photopic light but an unexpected excitation of scotopic rods and ON-bipolar cell responses (Figures 5A,B), indicating signs of retinal dysfunction. Compared with WT controls, homozygous Q84/Q84 mice showed both reduced photopic and scotopic a-wave and b-wave amplitudes, which indicated decreased function of, respectively, cones and rods in the outer photoreceptor layer as well as cells in the inner retina, predominantly Müller and bipolar cells (Figure 5A,B). Interestingly, the cone-mediated responses measured by the flicker test was nearly undetectable in Q84/Q84 mice (Figure 5B) suggesting depletion and/or highly dysfunctional cones in these retinas.

**Figure 5.**
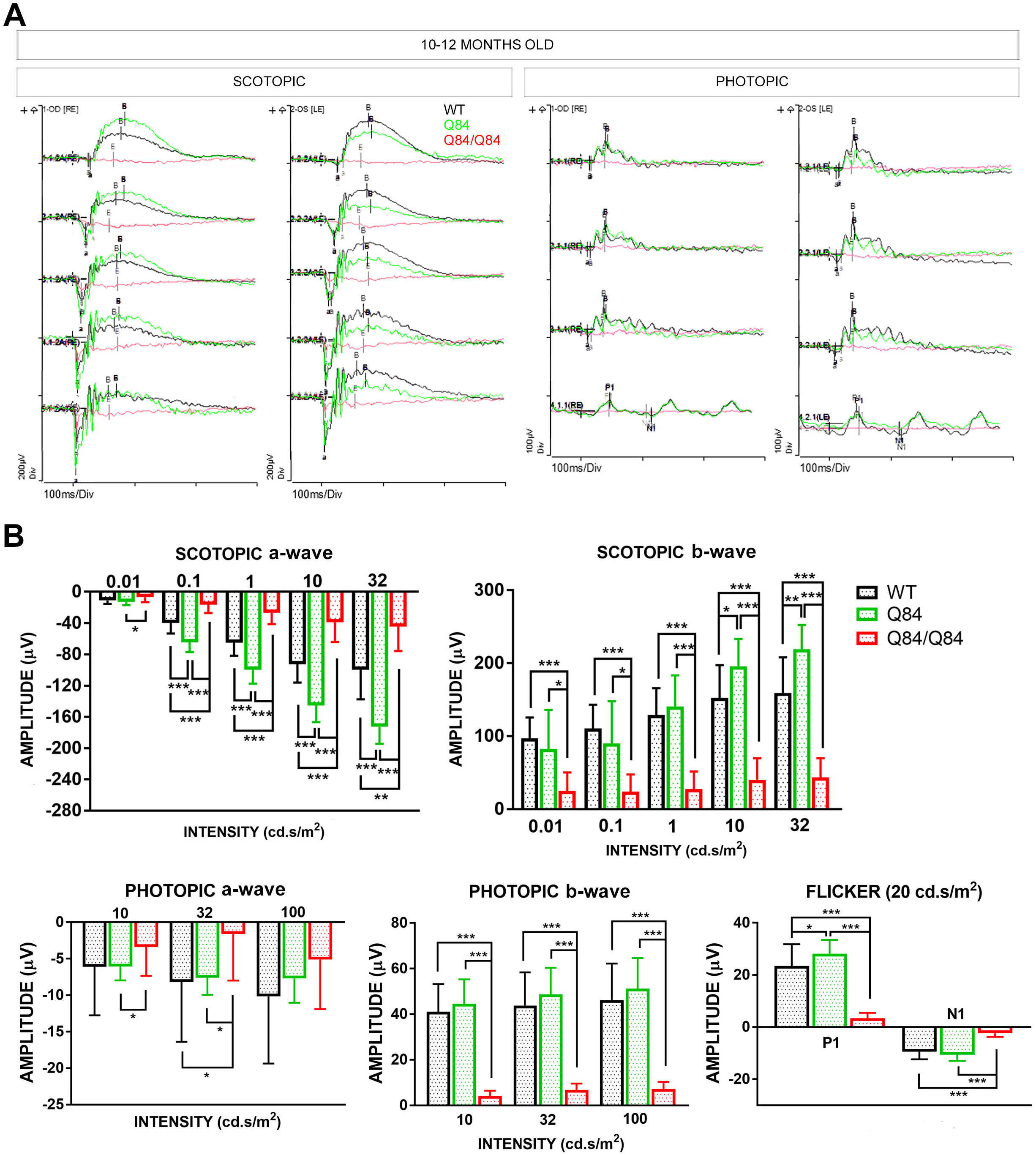
Retinal electrophysiological analyses reveal severe retinal dysfunction accompanied by highly decreased photoreceptor response in homozygous Q84/Q84 mice. **(A)** Representative ERG recordings of the scotopic (rod activity), photopic (cone activity) and flicker response of homozygous Q84/Q84 and hemizygous Q84, compared to WT mouse retinas (red, green and black, respectively) show severe loss of photoreceptor activation in response to light in Q84/Q84 mouse retinas. **(B)** Quantification of ERG recordings of scotopic (rods, a and b-wave), photopic (cones, a and b-waves) and flicker responses of hemizygous and homozygous Q84 retinas compared to WT (red, green and black, respectively) show nearly null photoreceptor response under all light conditions for homozygous Q84/Q84 mice. Scotopic response of femizygous Q84 retinas show hyperactivation of rods (a-wave) and ON-bipolar cells (b-wave). Statistical significance, * p<0.05, ** p<0.01, *** p<0.001, two eyes of 5-7 animals per group.

Briefly, all the electrophysiological tests confirmed altered function of rod photoreceptors in Q84 and Q84/Q84 mice, with the latter showing a severe phenotype including a drastic reduction of cone activity (Figure 5B). These electrophysiological alterations in mice reproduce in some extent the retinal dysfunction observed in patients with Spinocerebellar ataxia type 3 in this study.

### Expanded-polyQ ATXN3 progressively aggregates in several retinal layers of Spinocerebellar ataxia type 3 transgenic mice

Because several retinal cell types showed to be structurally altered and dysfunctional in Q84 and Q84/Q84 transgenic mice compared with WT controls and expanded-polyQ ATXN3 forms aggregates in brains of Spinocerebellar ataxia type 3 patients^11^ and in these transgenic mice^41,46^, we next analyzed whether ATXN3 also aggregates in retinal cells of Q84 and Q84/Q84 mice possibly impacting their function and health. Immunostaining of ATXN3 in retinal cryosections of 8-month-old and 1.5-year-old Q84, Q84/Q84 and WT littermate mice showed indeed accumulation of ATXN3-positive aggregates and large inclusions in several layers of Q84 and Q84/Q84 retinas (Figure 5). Hemizygous Q84 retinas displayed small aggregates of ATXN3 distributed throughout the inner segment (IS) of the photoreceptors and large ATXN3 inclusions in the GCL at 8 months-of-age (Figure 6A) and at 1.5 years-of-age (Figure 6B). Doubling the dose of the pathogenic human *ATXN3* transgene caused robust aggregation of ATXN3 in the IS of photoreceptors, inner nuclear layer (INL) and GCL as observed in the retinas of 8 month-old homozygous Q84/Q84 mice (Figure 6A). The accumulation of ATXN3 aggregates increased with age and was detectable throughout the whole retina at 1.5 years-of-age, being more prominent in the IS of photoreceptors and GCL of Q84/Q84 retinas (Figure 6B).

**Figure 6.**
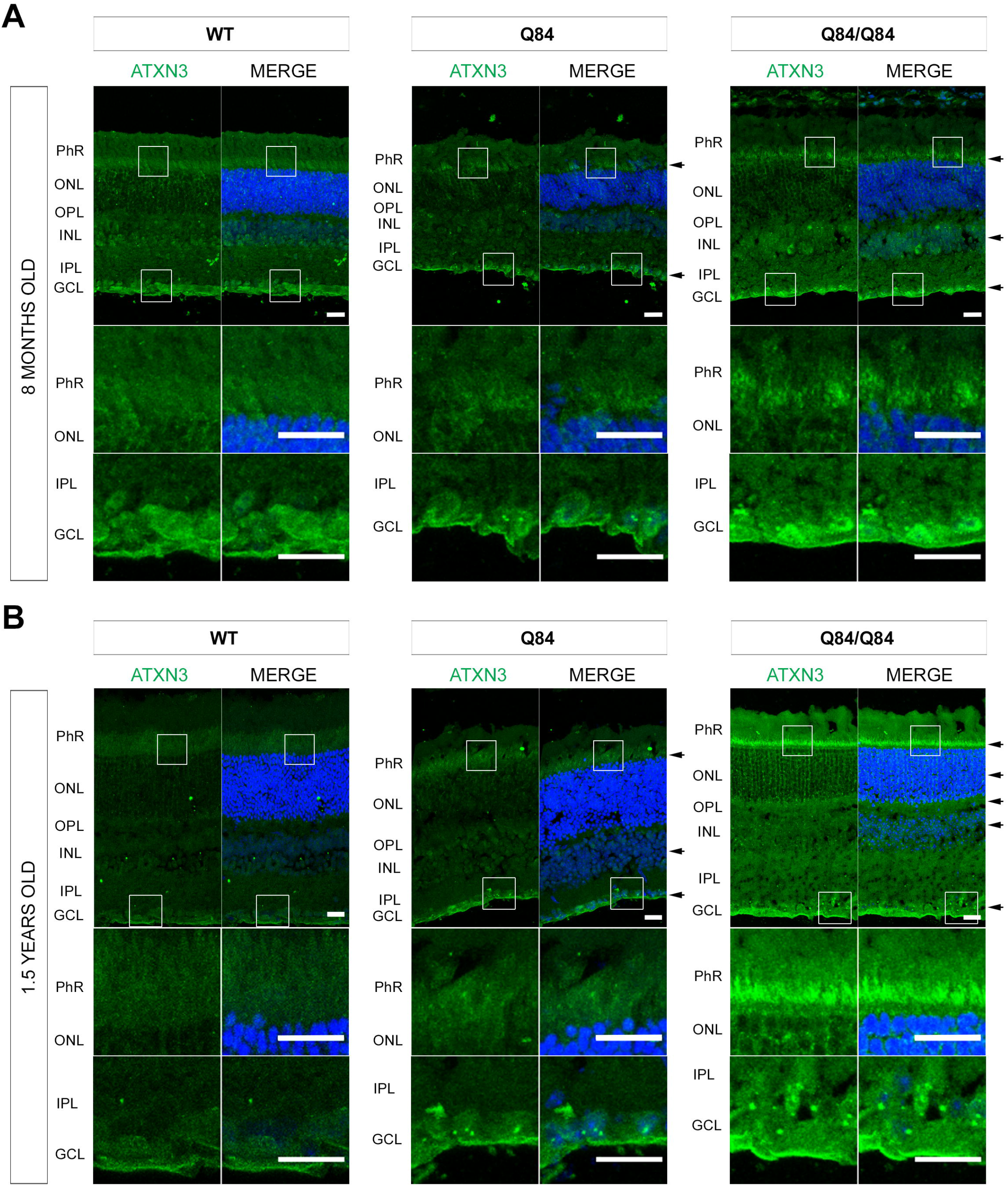
ATXN3 aggregates and forms large inclusions in the retinas of Q84 transgenic mice. Immunodetection of endogenous murine Atxn3 and human expanded ATXN3Q84 proteins in retinas from **(A)** 8 month-old and **(B)** 1.5 year-old mice shows ATXN3 inclusions and aggregates (in green) in the photoreceptor inner segments (IS), nuclear layers and ganglion cell layer (black arrows on the side of images) in the transgenic compared to the wild-type retinas. ATXN3 aggregates are at least detectable in 8 month-old mouse retinas but dramatically increase with aging, particularly in the IS of photoreceptors. Nuclei are counterstained with DAPI (blue). Magnification images below zoom into the photoreceptor and ganglion cell layers. Scale bar in all images corresponds to 20 μm. PhR - photoreceptors, ONL - outer nuclear layer, OPL - outer plexiform layer, INL - inner nuclear layer, IPL - inner plexiform layer, GCL - ganglion cell layer.

In short, ATXN3 preferentially aggregates in retinal layers that showed pronounced thinning by OCT in 10-12 month-old mice forming large inclusions in GCL of Q84 and Q84/Q84 mice, and small puncta in the IS of photoreceptors of Q84/Q84 mice.

### Spinocerebellar ataxia type 3 mouse retinas exhibit several ultrastructural abnormalities

Considering that certain layers of SCA3 mouse retinas are more prone to thinning and accumulation of ATXN3-positive aggregates, we next evaluated by transmission electron microscopy (TEM) whether the retinal morphology was affected by the expression of the human mutant *ATXN3* transgene. Ultrastructural imaging of all retinal layers from GCL to RPE of 1.5-year-old Q84/Q84 (N=3) and WT (N=3) littermate mice revealed various alterations of the cellular architecture in the transgenic mouse retinas compared to controls (Figure 7). Comparing with WT, retinal ganglion cells (RGCs) in the GCL of Q84/Q84 retinas showed reduced soma area and increased negative electron-dense puncta (possibly corresponding to ATXN3 aggregates) in nuclei and soma (Figure 7A). Higher number of negative electron-dense puncta were also observed in cell processes in the IPL mainly associated with the cytoskeletal filaments (Figure 7B), and in nuclei and soma of cell bodies localized in the INL (Figure 7C) of Q84/Q84 retinas compared to controls. The OPL of Q84/Q84 retinas was in average 20% thinner than controls, displaying some disorganization by showing incorporation of some mislocalized PhR nuclei (Figure 7D). In addition, the RPE layer and apical microvilli of Q84/Q84 retinas were, respectively, 40% thicker and 97% longer than controls (Figure 7E). While OCT measurements evidenced thinning of the ONL and PhR layers (Figure 4C) and TEM evaluation showed a trend to thinning of the IS layer (Supplementary Figure 2) in Q84/Q84 mice compared to controls, there were no apparent differences concerning the structure of photoreceptor nuclei, IS and stacking of the outer segment (OS) membranous discs (Supplementary Figure 2) between the two groups.

**Figure 7.**
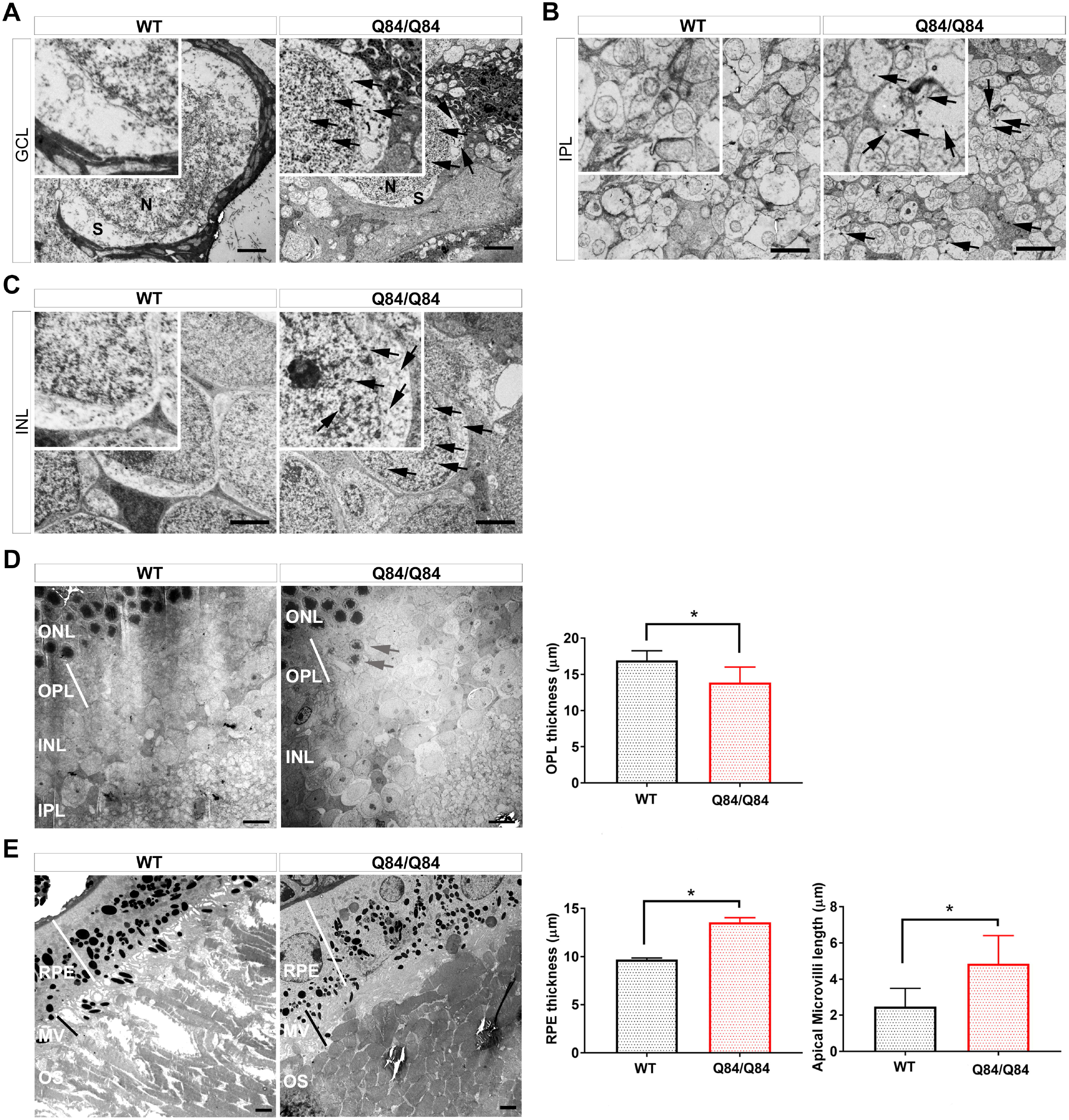
Spinocerebellar ataxia type 3 mouse retinas exhibit several ultrastructural abnormalities. Transmission electron microscopy (TEM) analysis reveals increased negative electron-dense puncta (black arrows) in **(A)** ganglion cell layer, **(B)** inner plexiform layer, and **(C)** inner nuclear layer of the Q84/Q84 retinas compared to controls. Scale bar corresponds to 2 μm. Square insets are magnifications of the zone of interest. The homozygous Q84/Q84 retinas show **(D)** thinner outer plexiform layer (scale bar corresponds to 8 μm) harboring mislocalized photoreceptor nuclei (grey arrows), and **(E)** wider RPE layer and elongated apical microvilli compared to WT (scale bar corresponds to 2 μm). Statistical significance by Mann Whitney, * p<0.05, n=3 per group). RPE - retinal pigment epithelium, MV - apical microvilli, OS - outer segment, ONL - outer nuclear layer, OPL - outer plexiform layer, INL - inner nuclear layer, IPL - inner plexiform layer, GCL - ganglion cell layer.

Ultrastructural analysis of aged Q84/Q84 retinas revealed several alterations in all retinal layers including the accumulation of intracellular electron-dense puncta in GCL, IPL and INL, disorganized and thinner OPL, wider RPE layer, and longer apical microvilli compared with the controls.

### Spinocerebellar ataxia type 3 mouse retinas show depletion of cones

Although photoreceptors of 1.5-year-old Q84/Q84 and WT mice appeared to be structurally similar (Supplementary Figure 2), Q84/Q84 retinas exhibited thinning of OPL, ONL and PhR layers (Figures 4 and 7) and decreased electrophysiological activity of rods and cones (Figure 5) suggesting depletion and/or dysfunction of photoreceptors. Therefore, we next assessed whether the overall morphology and number of rods and cones were specifically affected in Spinocerebellar ataxia type 3 transgenic mice by conducting immunostaining in retinal cryosections and whole mount retinas. In the vertebrate retina, the number of rod photoreceptors highly exceeds the number of cone photoreceptors, which represent about 3% of all photoreceptors in mice. Detection of rods in retinal cryosections using rhodopsin immunolabeling (localizing in the outer segment of rods) revealed very similar distribution patterns and number of rods in 8-month-old and 1.5-year-old Q84, Q84/Q84 and WT littermate mice (Figure 8A). Double-staining of cones using fluorescent peanut agglutinin (PNA, a lectin that specifically binds a sugar in glycoproteins of the cone membrane) and immunolabelling of L- and M-opsins in retinal cryosections of the same transgenic and control mice revealed progressive loss of cones in SCA3 compared to WT retinas (Figure 8B). While the number of cones was similar in 8-month-old Q84 and WT retinas, it was visibly lower in 1.5-year-old Q84 retinas compared to controls. On the other hand, and compared to controls, the number of cones was highly decreased in 8-month-old and more so in 1.5-year-old Q84/Q84 retina (Figure 8B), evidencing a transgene dosage effect. To further assess the dimension of cone loss in Spinocerebellar ataxia type 3 retinas, we conducted cone-specific PNA-staining in whole mount retinas of 4-8-month-old Q84/Q84 (n=5) and WT (n=5) mice and observed that the number of cones was, in fact, significantly decreased in Q84/Q84 retinas (about 10% less of control numbers) (Figure 8C).

**Figure 8.**
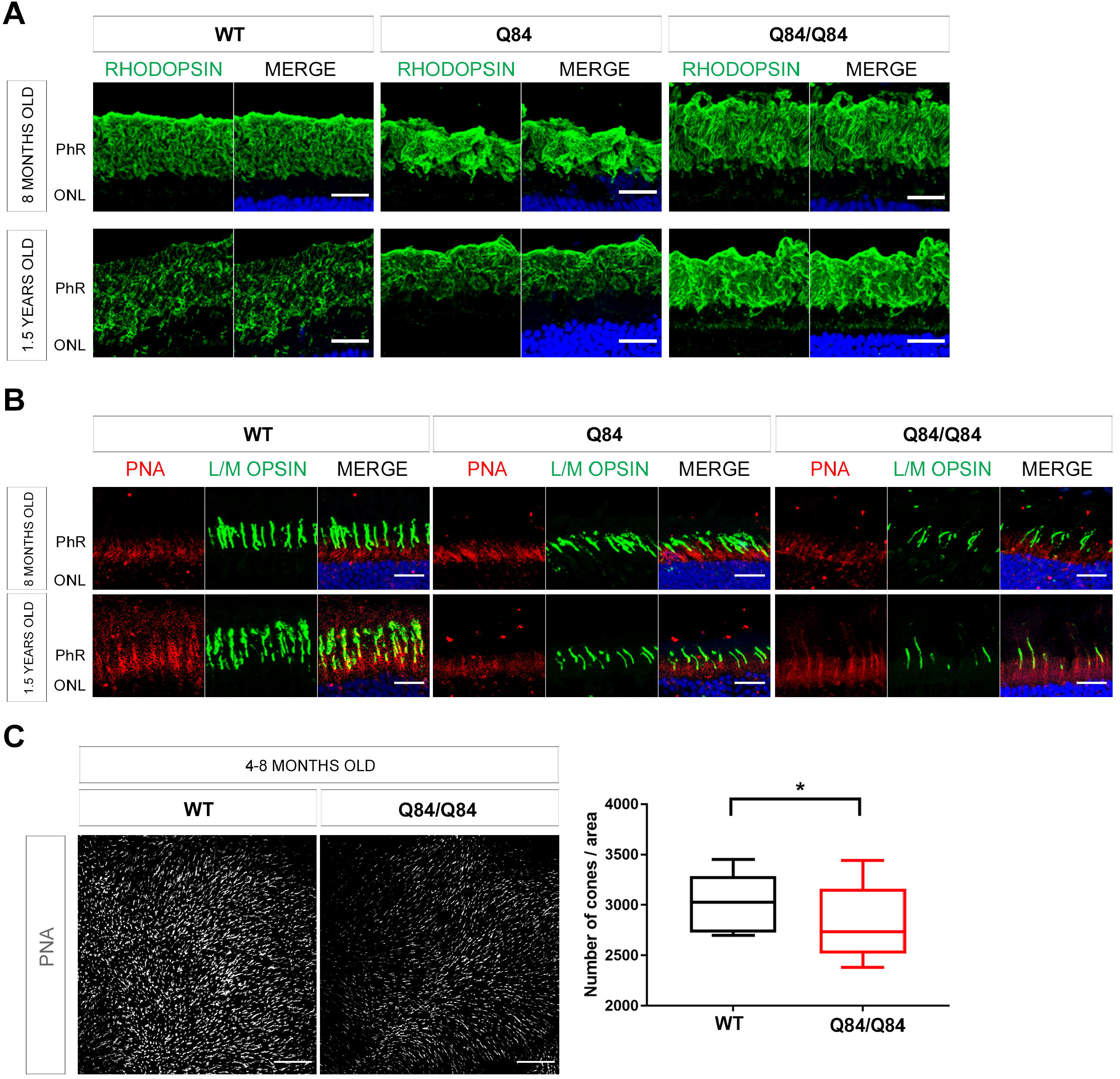
Expression of human pathogenic ATXN3Q84 in mouse retinas decreases the number of cone photoreceptor cells without other major photoreceptor alterations. **(A)** Immunohistochemistry of rhodopsin (green, labeling rods) in retinal cryosections (representative image) in 8 month-old and 1.5 year-old mice (representative images). Nuclei are counterstained with DAPI (blue). Scale bar 20 μm. Representative image of n=3 animals per group. **(B)** Immunohistochemistry of L/M opsin (green) and PNA (red, labeling cones) in retinal cryosections in 8 month-old and 1.5 year-old mice show that the number of cones decrease progressively with age correlating with the amount of expanded polyQ ATXN3 overexpression. Nuclei are counterstained with DAPI (blue). Scale bar 20 μm. **(C)** Whole mount retina staining show a statistically significant decrease in the total number of cones (stained with PNA, in white) in Q84/Q84 homozygous transgenic compared to wild-type mouse retinas (representative image) (n=5 per group, Two-way ANOVA Statistical Test, * indicate p<0.05). Scale bar 100 μm. PhR - photoreceptors, ONL - outer nuclear layer.

In brief, while the number of rods appeared to be preserved in Spinocerebellar ataxia type 3 transgenic mouse retinas, cone counts were progressively lower from around 4 months of age onwards in homozygous Q84/Q84 mice.

## DISCUSSION

Although the retina is part of the CNS, its privileged position represents “a window to the brain”, as some authors have proposed^24,53,54^. Cerebellum and the retina may share similar genetic networks and functional pathways, as shown by the shared interactome of ATXN7 (Spinocerebellar ataxia type 7) and CACNA1A (Spinocerebellar ataxia type 6) with EFEMP1 and FBLN5 involved in macular degeneration^55^. In fact, patients with Spinocerebellar ataxia type 7 show macular dysfunction and morphological retinal alterations^22,56,57^. Importantly, previous studies reported that patients with Spinocerebellar ataxia type 3 also reveal thinning of the NFL, GCL and macula^29–31,58^, but whether the retinal function is affected in patients with this disease remained uncovered. We, therefore, sought to evaluate both the anatomy and function of the retina in patients with Spinocerebellar ataxia type 3 and in a transgenic mouse model.

Our OCT evaluation of five patients with Spinocerebellar ataxia type 3 revealed mild to prominent thinning of the macula, NFL and GCL, and reduced macular volume, consistent with previous above-described findings^29–31^. Interestingly, we observed that disease duration negatively impacts the central macular thickness, and that disease severity (SARA) inversely correlates with the average macular thickness and macular volume, and GCL thickness. These architectural alterations were also confirmed by electrophysiological measurements by mfERGs, which detected specific dystrophic zones in the cone-rich macula but also in the peripheral ring of rods, and by ffERGs that detected clear alterations of the wave implicit time and amplitude in both scotopic and photopic conditions. Importantly, the electroretinograms revealed that the response of parafovea, rods and inner retinal cells deteriorates with the disease progression and that rods activity progressively declines with the disease severity. Although the results have been obtained in a small number of patients and need further confirmation, this accurate phenotypic analysis in human patients suggest that the degeneration of the central macula is mostly related to the time of disease onset and progression, whereas the more general affectation of all the macula and GCL is more related to disease severity. At least in this small cohort, females appeared to be more affected than males, but their comparative older age might be a relevant contributing factor. Future studies should be conducted to evaluate whether retinas of females are indeed more affected than retinas of males with Spinocerebellar ataxia type 3.

Notably, results in the humanized Spinocerebellar ataxia type 3 Q84 mouse fully confirm the impact of ATXN3 accumulation in the retina. Our integrative anatomical, morphological and functional retinal analyses of the model revealed that these mice replicate the progressive retinal alterations of the patients and further provided evidence of cone loss and several pathological and ultrastructural abnormalities. Progressive accumulation of expanded polyQ ATXN3 throughout several retinal layers is accompanied by retinal dysfunction and degeneration. We believe that the progressive neurological alteration of the retina observed in this study parallels the previously reported course of brain pathology and motor deficits in this model^46^. Interestingly, hemizygous Q84 retinas showed an intermediate functional phenotype and milder retinal thinning compared with homozygous Q84/Q84 mice, correlating with the increase of human mutant ATXN3Q84 expression and supporting that the phenotype observed in homozygous animals is solely due to the neurotoxicity of the overexpressed expanded polyQ ATXN3 protein. For instance, while the loss of cones is already detectable in 4-8 month-old Q84/Q84 retinas, this phenotype is only clearly observed in retinas of 1.5 year-old hemizygous Q84 mice. Nonetheless, cone loss alone cannot fully explain the nearly null electrophysiological recordings in Q84/Q84 mice in response to light. In this respect, we have previously shown in *Atxn3* knockout mice that ATXN3 regulates ciliary formation, intraciliary trafficking^33^. As we observed that one of the preferential retinal sites of expanded polyQ ATXN3 accumulation is the photoreceptor IS, we could speculate that some of the cytotoxicity of these aggregates is due to interference in the trafficking of relevant photoreception and phototransduction proteins into the membranous disks of the photoreceptor OS, a highly specialized ciliary structure. This trafficking interference would also support the observed null function of cones and highly reduced light response of rods in addition to photoreceptor attrition in homozygous Q84/Q84 retinas. Besides, and similarly to *Atxn3^-/-^* retinas, Q84/Q84 retinas showed longer apical microvilli^33^ and thicker RPE, thus suggesting possible deficiencies of photoreceptor OS phagocytosis that could potentially contribute to the observed photoreceptor dysfunction and viability (thinning of ONL and PhR). Our combined results in the two mouse models suggest that the expanded *ATXN3* CAG repeat confers a combination of gain-of-function (as detected by hyperexcitability of rods in Q84 hemizygotes) and loss-of function (in Q84 homozygotes, in which rapid accumulation of ATXN3 aggregates might impair ATXN3 normal function) in the retina. Future studies are, however, needed to clarify this hypothesis.

Concerning photoreceptors, rods and cones were not equally affected in Q84 mice. In old animals, rod structure seemed to be preserved, even though rod response to scotopic light was highly diminished. In contrast, hemizygous Q84 mouse retinas showed a similar cone response to the wild-type animals, whereas displaying increased rod response, potentially reflecting hyperexcitability of these photoreceptors. According to our data, either high overexpression of the expanded polyQ ATXN3 or an extended disease duration (associated to an increased accumulation of mutant polyQ protein aggregates in retinal cells) might be required for profound photoreceptor dysfunction leading to cell death. The latter hypothesis would be in agreement with our results in human patients, in which disease duration and severity were relevant factors for retinal layer thinning and electrophysiological dysfunction. Importantly, using OCTs and ERGs we also verified that the expansion of the *ATXN3* CAG repeat in humans and mice impacts the health and activity of ON-bipolar, Müller and ganglion cells, besides photoreceptors. Accurate imaging in transgenic mouse retinas further confirmed this phenotype by: 1) immunofluorescent detection of high amounts of toxic ATXN3-positive aggregates in the INL (formed by nuclei of bipolar and Müller cells, among others), in the OPL (formed by synapses of photoreceptors with bipolar and other cells as well as processes of Müller cells holding photoreceptors in place), and in RGCs; and 2) TEM imaging, which revealed OPL thinning and disorganization (further supporting dysfunction of Müller cells and of photoreceptor synapses with bipolar cells), increased electro-dense puncta (potentially corresponding to toxic ATXN3 aggregates) throughout the cell bodies in the INL, cell processes in the IPL, and the nuclei and soma of RGCs. Although many neuronal layers of the mouse retina are affected by the toxic accumulation of ATXN3 aggregates, the most affected cells from the morphological point of view are cone photoreceptors, since an early phenotypic trait is cone attrition, whereas altered electrophysiological records are detected for both rods and cones. In this context, the phenotypic alterations displayed by *Atxn3* knockout mouse retinas^33^ is again worth mentioning, since ablation of *Atxn3* affected ciliary OS structure of both rod and cones, but particularly protein ciliary trafficking in cones and cone electrophysiology was highly altered, positing *Atxn3* as a potential candidate for retinal/macular disorders.

Overall, our results in human patients with Spinocerebellar ataxia type 3, supported by the retinal phenotype observed in the humanized transgenic mice, suggest that several structural and functional retinal deficiencies could represent a biological marker of the disease progression and severity. We propose that these progressive retinal dysfunction, particularly macular alterations, may be a shared signature of polyQ spinocerebellar ataxias^20–23^ or even potentially of rare cerebellar diseases in general. Further studies of a larger cohort of patients and pre-symptomatic carriers of mutations causing Spinocerebellar ataxia type 3 would be required to better understand these specific pathogenic mechanisms occurring in their retinas, and discern whether macular and GCL thinning, and early cone/rod electrophysiological dysfunction are accurate indicative biomarkers of the onset and progression of these neurological disorders.

In conclusion, non-invasive eye examination could potentially help clinicians detecting early signs of neurological disorder in carriers of mutations that cause cerebellar ataxias and constitute a relevant marker for these diseases, whose early clinical diagnosis would enable the access to future effective therapies before severe neurodegeneration occurs.

## ACKNOWLEDGEMENTS

The authors would like to thank: Henry L. Paulson for providing the anti-MJD antibody; Pennelope Blakely Kunkle and Sasha Meshinchi (Microscopy Core, Michigan Medicine) for TEM assistance; and Cheng-mao Lin and Sarah Sheskey (Functional Assessment Core at the Kellogg Eye Center, Michigan Medicine) for assistance with the mouse OCT and ERG evaluations. We are also indebted to the patients for their consent in participating in this research.

## FUNDING

This research was supported by grants SAF2016-80937-R (financed by MCIN/ AEI /10.13039/501100011033/ FEDER “Una manera de hacer Europa”), PID2019-108578RB-I00 (financed by MCIN/ AEI / 10.13039/501100011033), 2017SGR-0738 (Generalitat de Catalunya), and La Marató TV3 (Project Marató 201417-30-31-32) to G.M.. V.T. was fellow of the MINECO (BES-2014-068639, Ministerio de Economía, Industria y Competitividad, Spain) and recipient of the grants EEBB-I-16-11823 and EEBB-I-17-12664 (Ministerio de Economía, Industria y Competitividad, Spain) for short international stages at the University of Michigan. This research was also supported by National Ataxia Foundation (SCA3 Translational Research Award 2019/2020) and University of Michigan discretionary funds to M.C.C..

## AUTHOR CONTRIBUTIONS

M.C.C. and G.M. designed and supervised the experimental work; M.C.C. and G.M. provided the funding; V.T. and M.C.C. performed the mouse experiments; R.C-M., A.C-C., M-F-R., performed the ophthalmological evaluation, OCT and ERGs tests in patients; B.S-D. and E.M. provided the clinical data of patients; N.S.A. managed mouse colonies, and assisted with mouse tissue collection; A.F.F. assisted with statistical analysis; N.K. contributed to the mouse ERG analysis; V.T, R.C-M., M.C.C., and G.M. wrote the manuscript; all the authors analysed the results, participated in the discussion and revised the manuscript.

## DECLARATION OF INTEREST

All the authors declare they have not competing interests.

## SUPPLEMENTARY MATERIAL

Supplementary material is available at *Brain* online.

## Abbreviations

BCVA: best-corrected visual acuity
ERG: electroretinogram
ffERG: full-field flash electroretinogram
GCL: ganglion cell layer
ICH: immunofluorescent histochemistry
IF: immunofluorescence
INL: inner nuclear layer
IPL: inner plexiform layer
IS: inner segment
mfERG: multifocal electroretinogram
MJD: Machado-Joseph disease
MV: apical microvilli
NFL: retinal nerve fiber layer
OCT: optical coherence tomography
ONL: outer nuclear layer
OPL: outer plexiform layer
OPs: oscillatory potentials
OS: outer segment
PhR: photoreceptor
PNA: peanut agglutinin
RAD: response activity density
RPE: retinal pigment epithelium
SARA: Scale for the Assessment and Rating of Ataxia
SCA3: Spinocerebellar ataxia type 3
SD-OCT: spectral domain-optical coherence tomography
TEM: transmission electron microscopy

## Notes

### Competing Interest Statement

The authors have declared no competing interest.

