## Supplementary Material for "Altered retinal structure and function in Spinocerebellar ataxia type 3"

### SUPLLEMENTARY FIGURE LEGENDS

**Supplementary Figure 1. Expression of human ATXN3 in the YACMJD84.2 (Q84) transgenic mouse retinas.**

Expression of transgenic human ATXN3(Q84) compared to the endogenous murine ATXN3 protein in retinas from hemizygous (Q84) and homozygous (Q84/Q84) transgenic mice and from wild-type (WT) littermate mice. Immunodetection of ATXN3 with the polyclonal anti-MJD antibody recognizing both human and mouse proteins confirmed the high levels of expression of the expanded human allele in retinas of the transgenic mice.

**Supplementary Figure 2. Transmission Electron Microscopy (TEM) of retinal ultrathin sections of homozygous Q84/Q84 and WT littermate mouse eyes.**

No gross morphological differences were apparent between the rod and cone photoreceptors from the homozygous transgenic and wild-type mouse retinas. Upper images show the similar length and morphology of the inner and outer segments of retinal photoreceptors in the two genotypes. Scale bar corresponds to 8 µm. Lower panels show the similarity in the connecting cilium length and membranous cone and rod disks structure in the two genotypes. Scale bar corresponds to 200 nm. Graphs show the average of IS and OS length ± STD of Q84/Q84 and WT retinas (n=3 animals from genotype). OS, outer segment; IS, inner segment; ONL, outer nuclear layer.

### Supplementary Figure 1

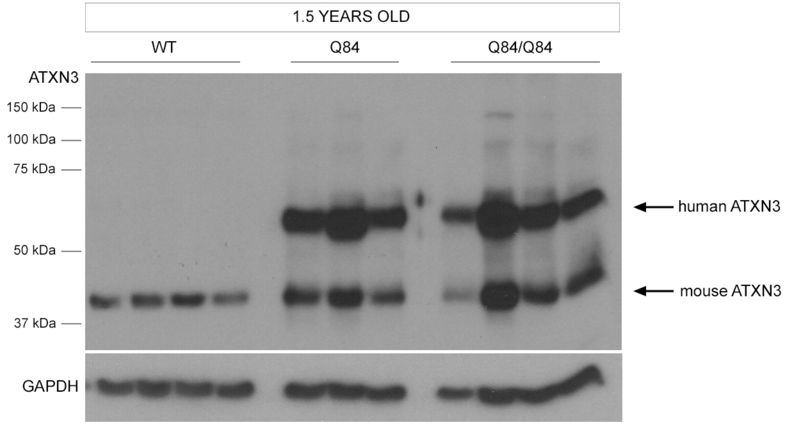

**Supplementary Figure 2**

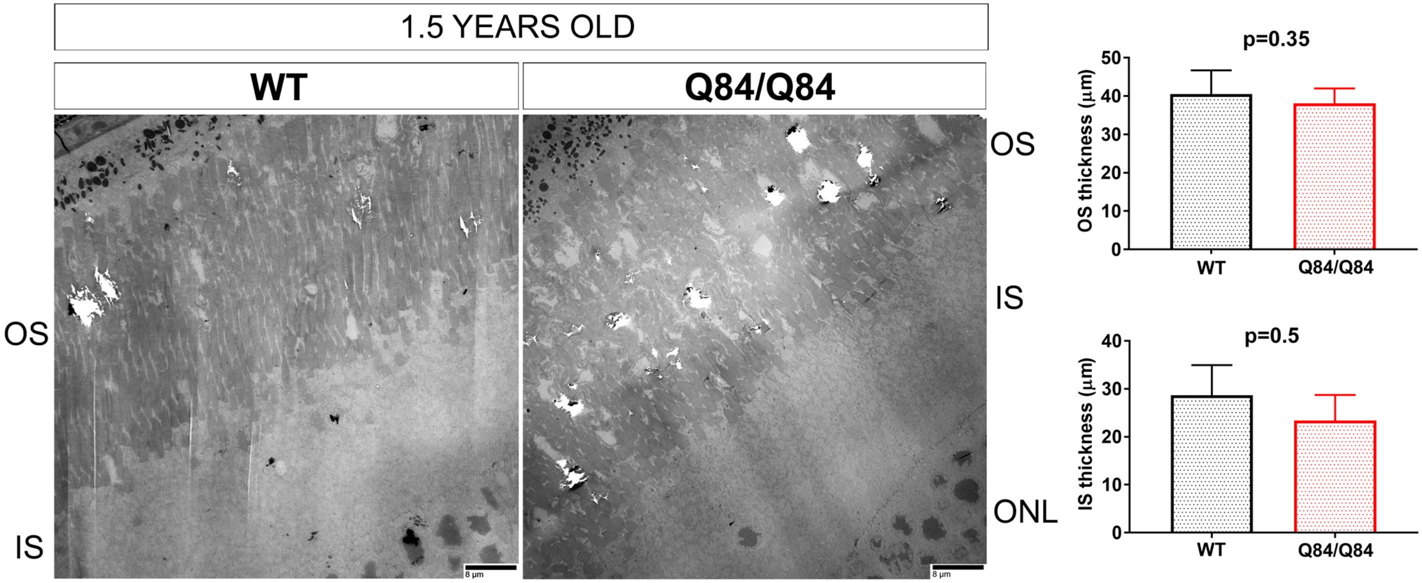


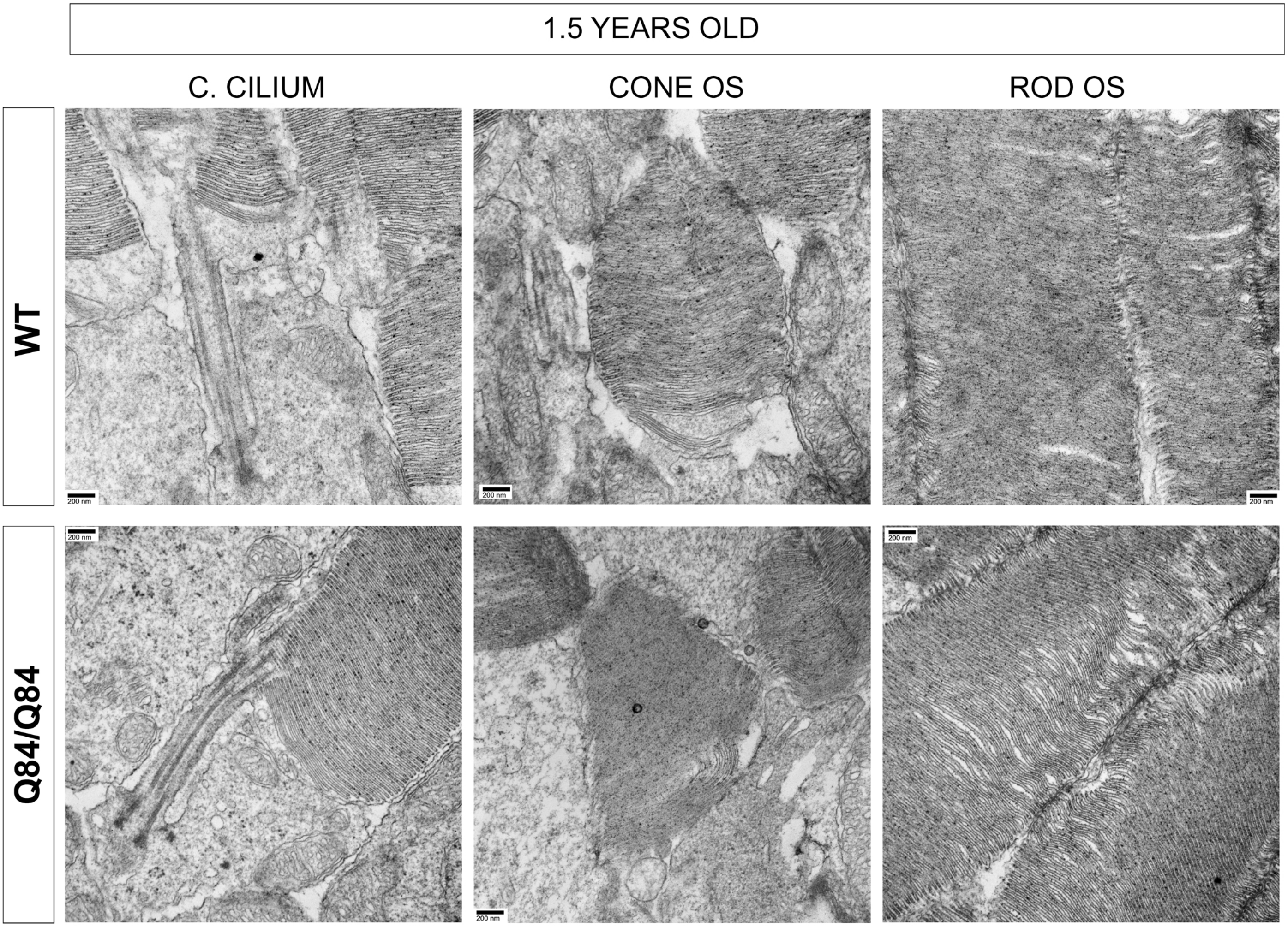

### Supplementary Table 1: Clinical and demographic data, OCT and ERG findings in patients with Spinocerebellar ataxia type 3 evaluated in this study

| | | | | | | | | | OCT (retina / optic disk) | | | | | | |
|---|---|---|---|---|---|---|---|---|---|---|---|---|---|---|---|
| | | | | | | | | | central | average | volume | GCL | NFL | NFL symmetry | comments |
| | age | gender | DD | SARA | DS | CAGs | eye | BCVA | µm | µm | mm$^3$ | µm | µm | % | |
| P1 | 54 | F | 10 | 6 | 1 | 68 | RE | 20/25 | 280 | 299 | 10.8 | 82 (78-85) | 88 (87-123) | 77 | **macular peripapillary thinning (BE), superior NFL peripapillary thinning (BE)** |
| | | | | | | | LE | 20/20 | 280 | 304 | 10.9 | 84 (82-91) | 102 (82-117) | | |
| P2 | 54 | M | 8 | 6.5 | 1 | 67 | RE | 20/20 | 284 | 280 | 10.1 | 78 (76-81) | 84 (51-108) | 92 | normal |
| | | | | | | | LE | 20/20 | 284 | 280 | 10.1 | 77 (76-80) | 80 (54-107) | | |
| P3 | 83 | F | 16 | 21.5 | 3 | 61 | RE | 20/25 | 227 | 261 | 9.3 | 72 (69-74) | 82 (51-104 | 88 | **temporal NFL peripapillary thinning (BE); mild GCL sectoral temporal thinning (BE)** |
| | | | | | | | LE | 20/25 | 226 | 258 | 9.2 | 71 (67-73) | 85 (49-114) | | |
| P4 | 62 | F | 23 | 19 | 3 | 70 | RE | 20/30 | 235 | 262 | 9.4 | 72 (69-75) | 85 (55-113) | 89 | **mild GCL sectoral temporal thinning (BE)** |
| | | | | | | | LE | 20/25 | 235 | 277 | 10.0 | 73 (71-75) | 86 (53-111) | | |
| P5 | 45 | M | 14 | 16 | 3 | 74 | RE | 20/20 | 282 | 263 | 9.5 | 68 (65-72) | 77 (60-106) | 92 | **macular peripapillary thinning (BE), inferior NFL peripapillary thinning (BE), GCL thinning (BE)** |
| | | | | | | | LE | 20/20 | 280 | 261 | 9.4 | 68 (65-71) | 74 (57-100) | | |

| | | mfERG | | | | | | | | | |
|---|---|---|---|---|---|---|---|---|---|---|---|
| | | R1 | | R2 | | R3 | | R4 | | R5 | |
| | eye | IT | RAD | IT | RAD | IT | RAD | IT | RAD | IT | RAD |
| P1 | RE | 48.0 | 95.2 | 42.2 | 47.1 | 45.1 | 25.5 | 41.2 | 11.5 | 40.2 | 7.1 |
| | LE | 47.1 | 72.2 | 50.0 | 43.8 | 41.2 | 19.5 | 45.1 | 9.8 | 43.1 | 8.9 |
| P2 | RE | 50.0 | 154.5 | 48.0 | 78.5 | 46.1 | 46.9 | 45.1 | 29.9 | 45.1 | 25.7 |
| | LE | 47.1 | 128.7 | 43.1 | 72.3 | 46.1 | 50.8 | 46.1 | 35.4 | 46.1 | 30.1 |
| P3 | RE | 50.0 | 78. | 47.1 | 31.0 | 49.0 | 23.9 | 47.1 | 12.0 | 45.1 | 9.1 |
| | LE | 54.9 | 60.0 | 47.1 | 35.8 | 45.1 | 11.7 | 44.1 | 11.2 | 45.1 | 7,4 |
| P4 | RE | 46.1 | 91.1 | 42.2 | 38.3 | 49.0 | 23.3 | 47.1 | 11.4 | 48.0 | 7.0 |
| | LE | 52.0 | 75.1 | 48.0 | 31.5 | 41.2 | 23.5 | 45.1 | 9.4 | 47.1 | 11.7 |
| P5 | RE | 46.1 | 126.5 | 42.2 | 62.1 | 45.1 | 47.1 | 44.1 | 32.2 | 45.1 | 24.1 |
| | LE | 41.2 | 93.4 | 41.2 | 45.4 | 45.1 | 27.7 | 44.1 | 17.8 | 44.1 | 12.1 |

| | | ffERG | | | | | | | | | | | | | | | | | |
|---|---|---|---|---|---|---|---|---|---|---|---|---|---|---|---|---|---|---|---|
| | | DA 0.01 | | DA 1.0 | | | | DA 10.0 | | | | LA 3.0 | | | | DA 3.0 OPs | | LA 30 Hz flicker | |
| | | b | | a | | b | | a | | b | | a | | b | | P2 | OS2 | P1 | N1-P1 |
| | eye | T | A | T | A | T | A | T | A | T | A | T | A | T | A | T | A | T | A |
| P1 | RE | 94.9 | 96.5 | 18.5 | 133.4 | 47.8 | 198.1 | 17.3 | 168.5 | 49.8 | 222.0 | 16.4 | 29.5 | 31.9 | 100.4 | 27.0 | 25.3 | 63.9 | 76.2 |
| | LE | 99.6 | 142.2 | 18.5 | 147.1 | 49.8 | 226.3 | 17.6 | 177.9 | 49.8 | 252.2 | 16.7 | 35.6 | 30.8 | 110.4 | 27.0 | 25.4 | 63.9 | 87.3 |
| P2 | RE | 77.6 | 234.9 | 23.7 | 334.4 | 49.8 | 493.1 | 17.9 | 386.0 | 46.6 | 471.5 | 17.0 | 49.3 | 32.8 | 278.4 | 26.1 | 101.2 | 64.7 | 164.0 |
| | LE | 88.8 | 222.9 | 23.7 | 222.8 | 49.8 | 340.1 | 17.6 | 262.3 | 46.0 | 307.0 | 16.7 | 24.9 | 32.8 | 296.1 | 25.8 | 77.4 | 65.6 | 155.9 |
| P3 | RE | 99.9 | 88.0 | 24.0 | 95.8 | 47.2 | 154.7 | 19.6 | 132.5 | 47.5 | 144.0 | 29.6 | 7.3 | 17.3 | 16.3 | 26.7 | 21.2 | 65.6 | 38.7 |
| | LE | 91.4 | 110.3 | 19.6 | 127.8 | 46.3 | 219.6 | 18.2 | 157.5 | 46.9 | 229.1 | 16.7 | 2.0 | 41.6 | 10.8 | 27.2 | 25.7 | 65.9 | 64.6 |
| P4 | RE | 99.9 | 82.9 | 19.0 | 110.0 | 45.1 | 163.4 | 18.2 | 153.2 | 45.1 | 177.9 | 17.9 | 21.8 | 31.6 | 76.4 | 28.1 | 31.7 | 62.4 | 54.1 |
| | LE | 99.9 | 102.9 | 19.3 | 148.8 | 45.4 | 215.5 | 19.0 | 184.6 | 46.0 | 225.0 | 18.5 | 23.2 | 31.3 | 81.3 | 27.8 | 34.5 | 62.1 | 64.2 |
| P5 | RE | 97.6 | 307.5 | 19.6 | 305.8 | 48.6 | 498.4 | 18.5 | 315.9 | 49.8 | 434.3 | 17.3 | 105.0 | 31.9 | 297.4 | 27.8 | 51.2 | 65.6 | 205.7 |
| | LE | 99.9 | 212.9 | 18.5 | 264.9 | 49.8 | 390.2 | 17.9 | 317.4 | 49.8 | 385.8 | 16.4 | 72.8 | 31.9 | 292.2 | 26.7 | 41.3 | 65.0 | 178.1 |

DD, Disease duration, years from onset to the evaluation time; F, female; M, male; SARA, Scale for the Assessment and Rating of Ataxia; DS, disease stage; CAGs, number of CAG repeats; RE, right eye; LE, left eye; BCVA, best corrected visual acuity (Snellen chart); OCT, optical coherence tomography; central, foveal thickness; average, macular thickness; volume, of macular region; GCL, ganglion cell layer thickness; NFL, nerve fiber layer thickness; NFL symmetry between NFL; RE, right eye; LE, left eye; BE, both eyes; (minimum-maximum) thickness according to OSCAR-IB and APOSTEL[38,39]; ERG, electroretinogram; mfERG, multifocal ERG; R1-R5, Ring 1 – Ring 5; IT, N1/P1 implicit time (ms); RAD, response amplitude density (nV/deg2); ffERG, full-field flash ERG; DA, dark-adapted; LA, light-adapted; OP, oscillatory potentials; T, latency (ms); A, amplitude (nV); a, a-wave; b, b-wave
